# GCN5L1-mediated acetylation prevents Rictor degradation in cardiac cells after hypoxic stress

**DOI:** 10.1101/2023.10.26.564170

**Authors:** Paramesha Bugga, Janet R. Manning, Bellina A.S. Mushala, Michael W. Stoner, John Sembrat, Iain Scott

## Abstract

Cardiomyocyte apoptosis and cardiac fibrosis are the leading causes of mortality in patients with ischemic heart disease. As such, these processes represent potential therapeutic targets to treat heart failure resulting from ischemic insult. We previously demonstrated that the mitochondrial acetyltransferase protein GCN5L1 regulates cardiomyocyte cytoprotective signaling in ischemia-reperfusion injury *in vivo* and hypoxia-reoxygenation injury *in vitro*. The current study investigated the mechanism underlying GCN5L1-mediated regulation of the Akt/mTORC2 cardioprotective signaling pathway. Rictor protein levels in cardiac tissues from human ischemic heart disease patients were significantly decreased relative to non-ischemic controls. Rictor protein levels were similarly decreased in cardiac AC16 cells following hypoxic stress, while mRNA levels remained unchanged. The reduction in Rictor protein levels after hypoxia was enhanced by the knockdown of GCN5L1, and was blocked by GCN5L1 overexpression. These findings correlated with changes in Rictor lysine acetylation, which were mediated by GCN5L1 acetyltransferase activity. Rictor degradation was regulated by proteasomal activity, which was antagonized by increased Rictor acetylation. Finally, we found that GCN5L1 knockdown restricted cytoprotective Akt signaling, in conjunction with decreased mTOR abundance and activity. In summary, these studies suggest that GCN5L1 promotes cardioprotective Akt/mTORC2 signaling by maintaining Rictor protein levels through enhanced lysine acetylation.

## Introduction

Ischemic heart disease is the leading cause of death worldwide (Ferreira-Gonzalez, 2014). Preventive strategies, such as lifestyle modifications, weight management, blood pressure control, and lipid-lowering medications, are used to reduce ischemic heart disease-associated mortality (Virani et al., 2021). Furthermore, in acute myocardial infarction (AMI), rapid blood flow restoration and post-infarct treatment supported by dual antiplatelet medication, b-blockers, and angiotensin-converting enzyme (ACE) inhibitors lead to improved outcomes (Elgendy et al., 2019). Cardiomyocyte apoptosis and cardiac fibrosis are the primary reasons for morbidity among ischemic heart disease patients (Saraste et al., 1997), and ameliorating cardiomyocyte apoptosis is a promising therapeutic target for treating ischemic heart disease (Tripathi et al., 2023).

Rictor is an essential protein of the mTORC2 complex which regulates the Akt/mTORC2 cytoprotective signaling pathway (Sciarretta et al., 2015, Kaldirim et al., 2022, Volkers et al., 2013). The role of Rictor in cytoprotective signaling has been extensively studied in various cancers, metabolic diseases, and inflammatory diseases (Gruninger et al., 2022, Jebali et al., 2021, Oneyama et al., 2013, Kocalis et al., 2014). Our group previously reported that lysine acetylation of Rictor (Ac-Rictor) is essential to ameliorate loss of cardiomyocytes and cardiac cells in response to hypoxic injury (Manning et al., 2019b). However, the mechanism underlying the regulation of Rictor acetylation, and its subsequent role in cytoprotective signaling pathways, remained unknown.

GCN5L1 is ubiquitously present in mitochondria, and is involved in the regulation of mitochondrial protein acetylation (Scott et al., 2018, Thapa et al., 2020, Zhang et al., 2023a, Webster et al., 2013). Our group recently reported that GCN5L1 promotes cardiac mitochondrial protein acetylation (e.g. fuel substrate metabolism enzymes) and promotes fatty acid oxidation (Thapa et al., 2018, Manning et al., 2022). Though GCN5L1 is best known for its mitochondrial acetyltransferase activity, recent work has identified several acetylation substrates of GCN5L1 in the cytosol, including Rictor (Wu et al., 2018, Manning et al., 2019a). In our current study, we show for the first time that GCN5L1 is essential to maintain cytoprotective Akt/mTORC2 cardioprotective signaling upon hypoxic stress by preventing Rictor degradation.

## Materials and Methods

### Clinical Samples

Fresh human cardiac tissue samples were collected from the left ventricles of organ donors deemed not suitable for transplant, under a protocol approved by the University of Pittsburgh Committee for Oversight of Research and Clinical Training (CORID). Tissues were flash-frozen and stored at −80°C until processing. Non-ischemic: 2M/2F, age range 56-68 years, mean 60.25 years. Ischemic: 2M/2F, age range 49-88, mean 61.75 years.

### Cell Lines

AC16 cells were purchased from Sigma-Millipore (Cat No: SCC109). AC16 cells were created by the fusion of primary adult ventricular cardiomyocytes with SV40 fibroblasts, and mimic the protein expression and metabolic properties of primary human cardiomyocytes (Davidson et al., 2005, Li et al., 2019, Truong et al., 2015). AC16 cells were cultured under standard conditions (i.e., 25 mM glucose DMEM, 10% FBS, antibiotic-antimycotic, 5% CO_2_, at 37 °C; all ThermoFisher, USA).

### Generation of GCN5L1 knockdown and overexpressed stable cell lines

AC16 cells were transduced with shRNA and ORF lentiviral particles at a multiplicity of infection (MOI) of 10, with scrambled control or GCN5L1 shRNA (Sigma-Aldrich, USA), or control or GCN5L1 ORF lentiviral particles (Origene, USA), followed by puromycin selection. RT-qPCR and western blotting confirmed GCN5L1 knockdown and overexpression.

### Measurement of Rictor mRNA Translation and Degradation

Rictor protein stability was measured in control and genetically modified (GCN5L1 knockdown or overexpressed) AC16 cells by challenging them with cycloheximide, a protein synthesis inhibitor, and MG132, a proteosome degradation inhibitor, at a concentration of 30 ng/ml or 0.5 μM, respectively, for 24 h.

### Hypoxia studies

After 24 h of culturing in 10 cm cell culture plates, hypoxia studies were performed on control, GCN5L1 knockdown, or GCN5L1 overexpressed AC16 cells. Cells were washed three times with 1X PBS, then incubated with Esumi Buffer (12 mM KCl, 0.9 mM CaCl_2_, 0.5 mM MgCl_2_, 137 mM NaCl, 20 mM HEPES, 20 mM 2-deoxy-D-glucose (2-DG), 20 mM lactic acid, pH 6.2) for 3 h (Chen and Vunjak-Novakovic, 2018, Yang et al., 2018). After completion of hypoxia treatment, cells were washed with 1X PBS and lysed in CHAPS lysis buffer, with added protease, phosphatase, and deacetylase inhibitors.

### RT-qPCR

RNA was harvested from cells using the RNEasy kit (Qiagen), with a μLITE analyzer (BioDrop, USA) used to evaluate the quality and quantity. RNA (1 μg) was used to synthesize cDNA using the Maxima Reverse Transcriptase (ThermoFisher, USA) kit, then gene expression studies were performed with specific primers for *Gcn5L1* (Cat.No. QT00016002), *Rictor* (Cat.No. QT00065793), and *GAPDH* (Cat.No. QT00079247; all Qiagen, USA).

### Immunoblotting

Cells and cardiac tissues were lysed in 1% CHAPS buffer. Protein samples were quantified using a μLITE analyzer (BioDrop), and 20 μg of proteins was separated on an SDS-PAGE gel. Protein bands were transferred onto 0.2/0.45 μm nitrocellulose membranes, which were blocked using Odyssey blocking buffer (Li-Cor, USA). Membranes were incubated with primary antibodies (GAPDH, Cell Signaling, Cat. 97166 [D4C6R], mouse mAb, 1:1000; Akt, Cell Signaling, Cat. 4685 [11E7], rabbit mAb, 1:1000; p-Akt [Ser473], Cell Signaling, Cat. 9271, Rictor, Cell Signaling, Cat. 2114 [53A2], rabbit mAb 1:1000; mTOR, Cell signaling, Cat.no. 2983 rabbit mAb 1:1000; p-mTOR cell signaling, Cat.no. 5536, rabbit mAb 1:1000; GCN5L1 1:500, rabbit pAb) overnight at 4 °C, washed 3X with PBST, and incubated at room temperature with fluorescent secondary antibodies for 1 h (800 nm anti-rabbit, 700 nm anti-mouse; Li-Cor). Protein bands were imaged using an Odyssey Imager (Li-Cor), and quantitated using Image-J software (NIH, USA).

### Immunoprecipitation

For Ac-Rictor measurements, AC16 cell lysates were prepared in 1% CHAPS in the presence of 100 μM Trichostatin-A and 5 mM nicotinamide to block deacetylase activity. 1 μg/μl (500 μg protein) samples were prepared and incubated overnight at 4 °C in primary antibody against acetyl-lysine (Cell Signaling; Cat No. 9814 (1:100)). Lysates were then incubated in a 1:1 mixture of Protein A and Protein G beads for 4 h at 4 °C, and washed thoroughly in PBS. Bound proteins were eluted into reducing SDS lysis buffer, followed by immunoblotting as described above.

### Statistical Analysis

Statistical analyses were performed using GraphPad Prism 7. Student’s t-tests were used for simple comparisons between two groups. A *P* value < 0.05 was considered significant. Sample numbers are listed in the figure legends. All data are represented as mean ± SEM.

## Results

### Rictor protein levels were significantly reduced in cardiac tissues from human ischemic heart disease patients, and in cardiac AC16 cells under hypoxia stress

Rictor is an essential component of the cytoprotective mTORC2 complex, and its mRNA and protein levels are dynamically regulated in various disease conditions (Oneyama et al., 2013, Zhao et al., 2014, Zhang et al., 2023b). In addition, Rictor protein levels are dynamically regulated under various stress conditions (e.g. ER stress, nutrient starvation, ischemic injury, etc.) (Park et al., 2020, Maejima, 2020). As cardiomyocyte protein levels are generally susceptible to stressful conditions, especially ischemic stress (Ghosh et al., 2020), we examined Rictor protein levels in cardiac tissues from ischemic heart disease patients and nonfailing heart tissue samples. We observed a substantial decrease in Rictor protein abundance in ischemic heart disease relative to non-failing controls (Fig. 1 A-B). As chronic ischemic disease may impact protein levels through several mechanisms, we tested whether Rictor was regulated by acute ischemic stress. We found a decrease in Rictor protein levels in cardiac AC16 cells following acute hypoxia that occurred in the absence of a significant change in mRNA levels (Fig. 1C-D and Fig. S1). These results were consistent with our patient data, and suggest that Rictor levels are rapidly regulated by ischemia.

**Figure 1.**
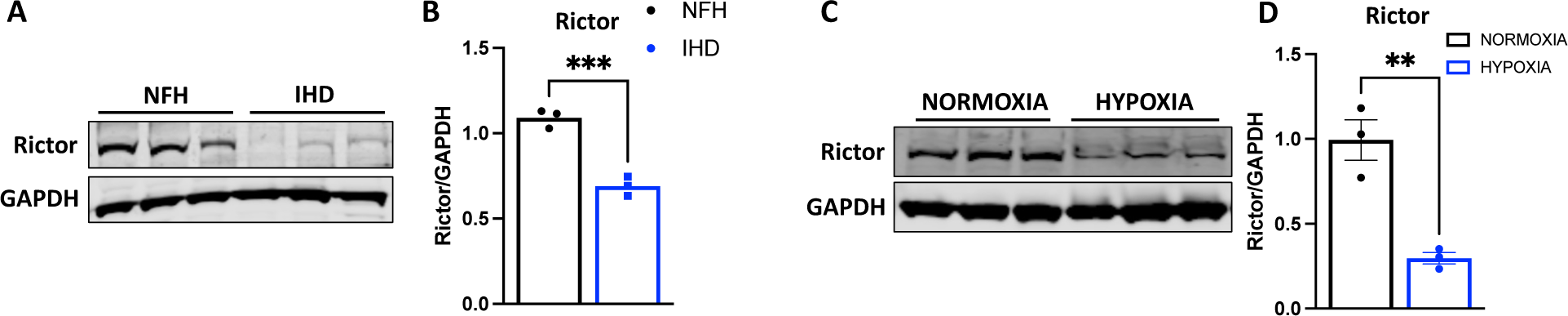
Rictor protein levels are substantially decreased in ischemic heart disease patient cardiac tissues and in AC16 cells after hypoxia. **A-B** Rictor protein levels in non-failing (NFH) and ischemic (IHD) cardiac tissues. **C-D** Rictor protein expression levels of AC16 cells in normoxia and *in vitro* hypoxia conditions. Data were expressed as Mean±SEM, n=3, **p>0.01 vs Non-failing heart samples, ***p>0.001 vs Normoxia treated AC16 cells.

### Genetic manipulation of GCN5L1 regulates Rictor protein abundance in the absence of transcriptional changes

GCN5L1 is well known for its mitochondrial acetyltransferase activity, which regulates cellular energy metabolism through its acetylation of various metabolic enzymes (e.g. those involved in fatty acid oxidation and pyruvate oxidation)(Thapa et al., 2020, Manning et al., 2019a). We recently reported that GCN5L1 also impacts the acetylation status of cytosolic proteins, including Rictor (Manning et al., 2019a). To study the role of GCN5L1 on Rictor protein acetylation and its cytoprotective signaling, we generated GCN5L1 knockdown (KD) and overexpressed (OE) stable AC16 cell lines using lentiviral particles expressing either GCN5L1-targeted shRNA or the GCN5L1 ORF.

We measured Rictor protein and mRNA levels in GCN5L1 KD and OE AC16 cells at baseline, and found a significant decrease in Rictor protein in KD cells, and a significant increase in Rictor protein levels in the overexpression condition (Fig. 2 A,C and Fig. 3 D,E). Surprisingly, we did not see any changes in mRNA levels in both conditions (Fig. 2 E and Fig. 3 F). As these results were similar to our acute hypoxia study (i.e. decreased Rictor protein levels and no change in Rictor mRNA levels; Fig. 1C-D and Fig. S1), we examined how changes in GCN5L1 abundance may impact Rictor at the post-transcriptional level. We hypothesized that changes in GCN5L1-mediated Rictor acetylation may regulate its abundance. As such, we measured Ac-Rictor levels in control, GCN5L1 KD, and GCN5L1 OE AC16 cells. We found a significant decrease in Rictor protein acetylation in GCN5L1 KD cells, and increased Ac-Rictor levels in GCN5L1 OE cells (Fig. 2 F-G and Fig. 3 G-H). From these observations, we conclude that GCN5L1 regulates Rictor acetylation, and that this process is involved in the regulation of Rictor protein abundance.

**Figure 2.**
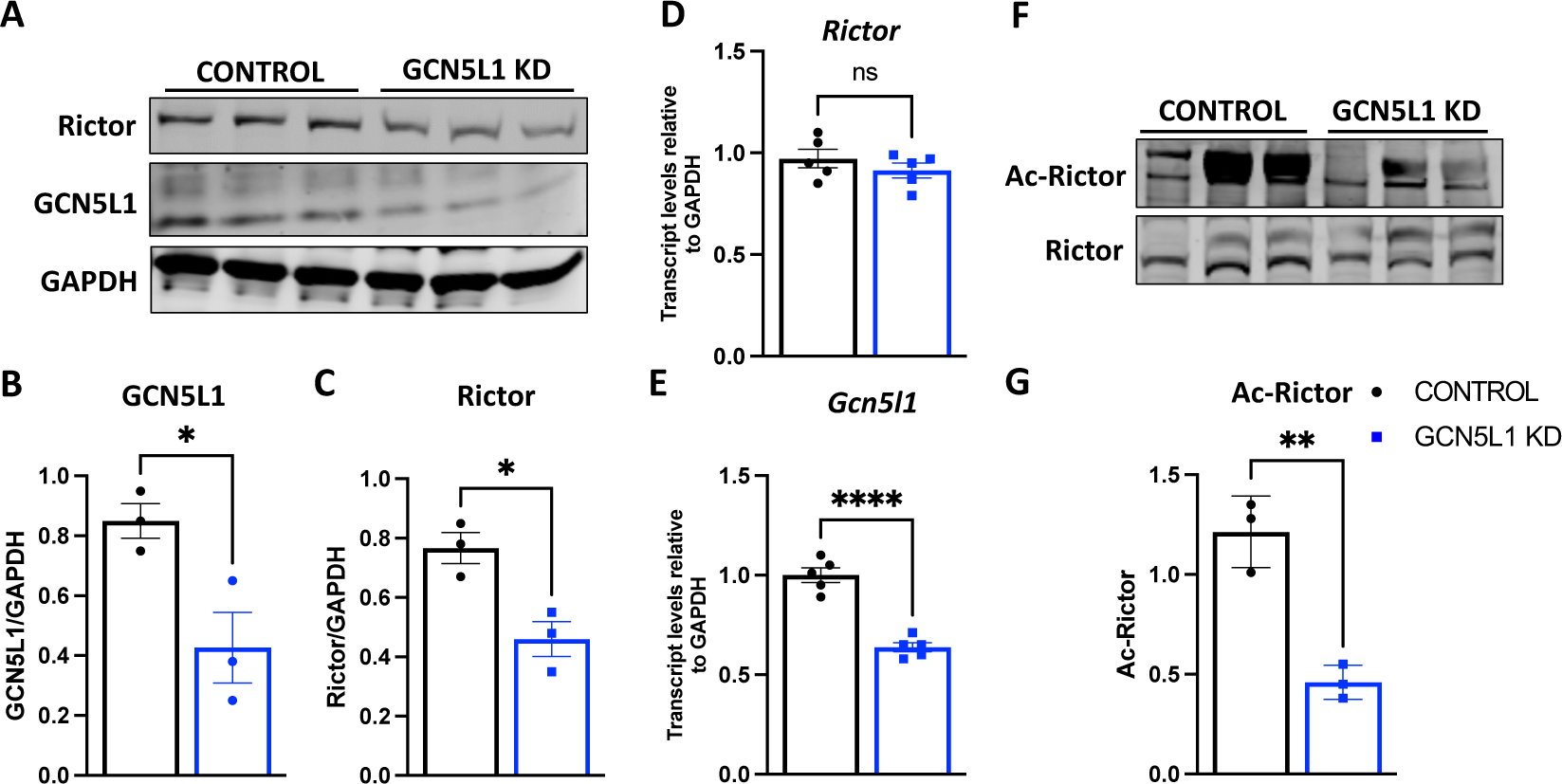
GCN5L1 knockdown in AC16 cells reduces Rictor protein abundance in the absence of gene expression changes. **A-C** GCN5L1 and Rictor Protein levels in control and GCN5L1 KD AC16 cells under basal conditions**. D** GCN5L1 mRNA expression levels in control and AC16 cells. **E** Rictor mRNA expression levels in control and GCN5L1 KD cells. **F-G** Ac-Rictor levels in control and GCN5L1 KD cells. Data were expressed as Mean±SEM, n=3, *p>0.05 vs Control AC16 cells **p>0.01 vs Control AC16 cells, ***p>0.001 vs Control AC16 cells.

**Figure 3.**
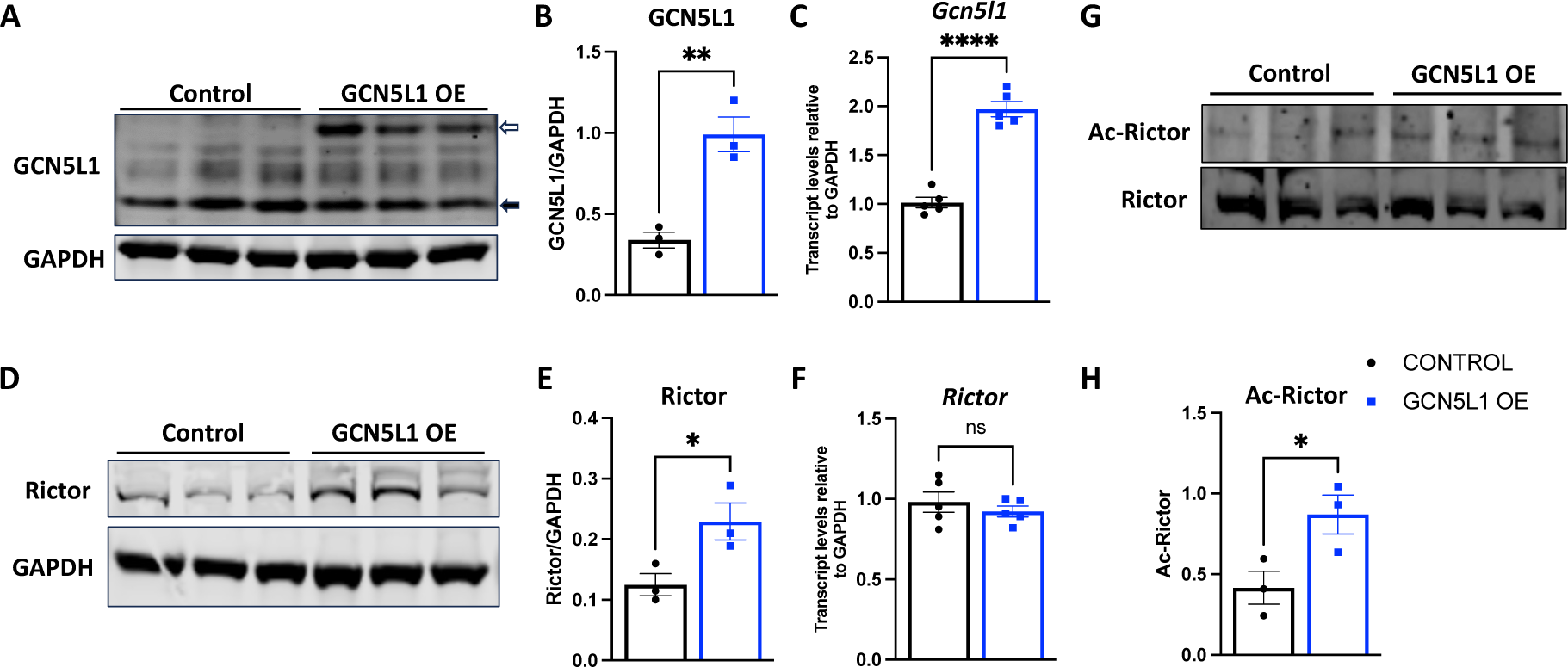
GCN5L1 overexpression increases the Rictor protein levels without influencing mRNA expression. **A-B** GCN5L1 protein levels in control and GCN5L1 OE AC16 cells in basal condition. Black arrow = endogenous GCN5L1; white arrow = GCN5L1 OE. **C-D** Rictor Protein levels in control and GCN5L1 OE AC16 cells in basal condition. **E** GCN5L1 mRNA expression levels in control and AC16 cells. **F** Rictor mRNA expression levels in control and GCN5L1 KD cells**. G-H** Ac-Rictor levels in control and GCn5L1 KD cells. Data were expressed as Mean±SEM, n=3, *p>0.05 vs Control AC16 cells **p>0.01 vs Control AC16 cells, ***p>0.001 vs Control AC16 cells.

### GCN5L1 prevents Rictor degradation by inhibiting the proteasomal degradation pathway

Several important mechanisms regulate protein levels, including gene transcription, mRNA translational activity, and protein degradation rates. We wanted to investigate further how Rictor protein levels were dynamically regulated in AC16 cells, and determine whether Rictor mRNA translational activity or Rictor protein degradation pathways were the primary regulatory mechanism. To study this, we treated AC16 cells with cycloheximide (a protein synthesis inhibitor that blocks mRNA translation) and MG132 (an inhibitor of protein degradation through the proteasomal degradation pathway). We found a moderate, non-significant decrease in Rictor protein levels when AC16 cells were treated with cycloheximide, but observed a substantial increase in Rictor protein levels when the proteasomal protein degradation pathway was inhibited by MG132 (Fig 4 A-D). We then measured Rictor protein levels in GCN5L1 KD and OE AC16 cells after treating them with cycloheximide and MG132. We observed a significant decrease in Rictor levels in GCN5L1 KD cells after cycloheximide treatment, and effect that was completely blocked in GCN5L1 OE cells under the same conditions. Similarly, the addition of MG132 to GCN5L1 KD cells led to a rapid accumulation of Rictor which, while remaining significant, was attenuated in GCN5L1 OE cells. Combined, we conclude that degradation of Rictor by the proteasome is the primary mechanism of Rictor protein regulation in cardiac cells. Furthermore, our data suggest that GCN5L1 acetyltransferase activity is a key mechanism which stabilizes Rictor protein and blocks it from proteasomal degradation.

**Figure 4.**
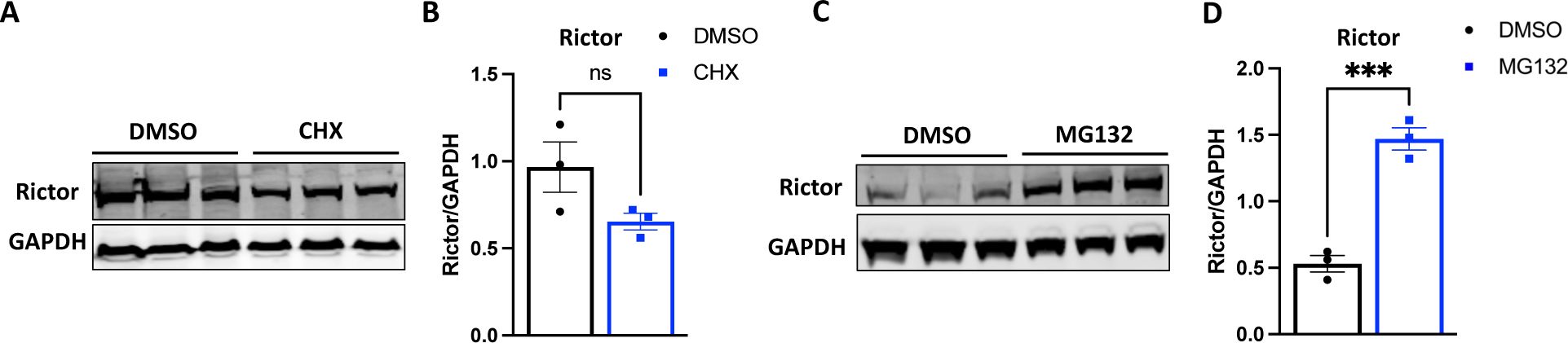
Rictor protein levels are regulated at the post-transcriptional level in AC16 cells. **A-B** Rictor protein levels in AC16 cells after challenging with Cycloheximide (CHX). **C-D** Rictor protein levels in control AC16 cells after MG132 treatment. Data were expressed as Mean±SEM, n=3, *p>0.05 vs DMSO **p>0.01 vs DMSO.

**Figure 5.**
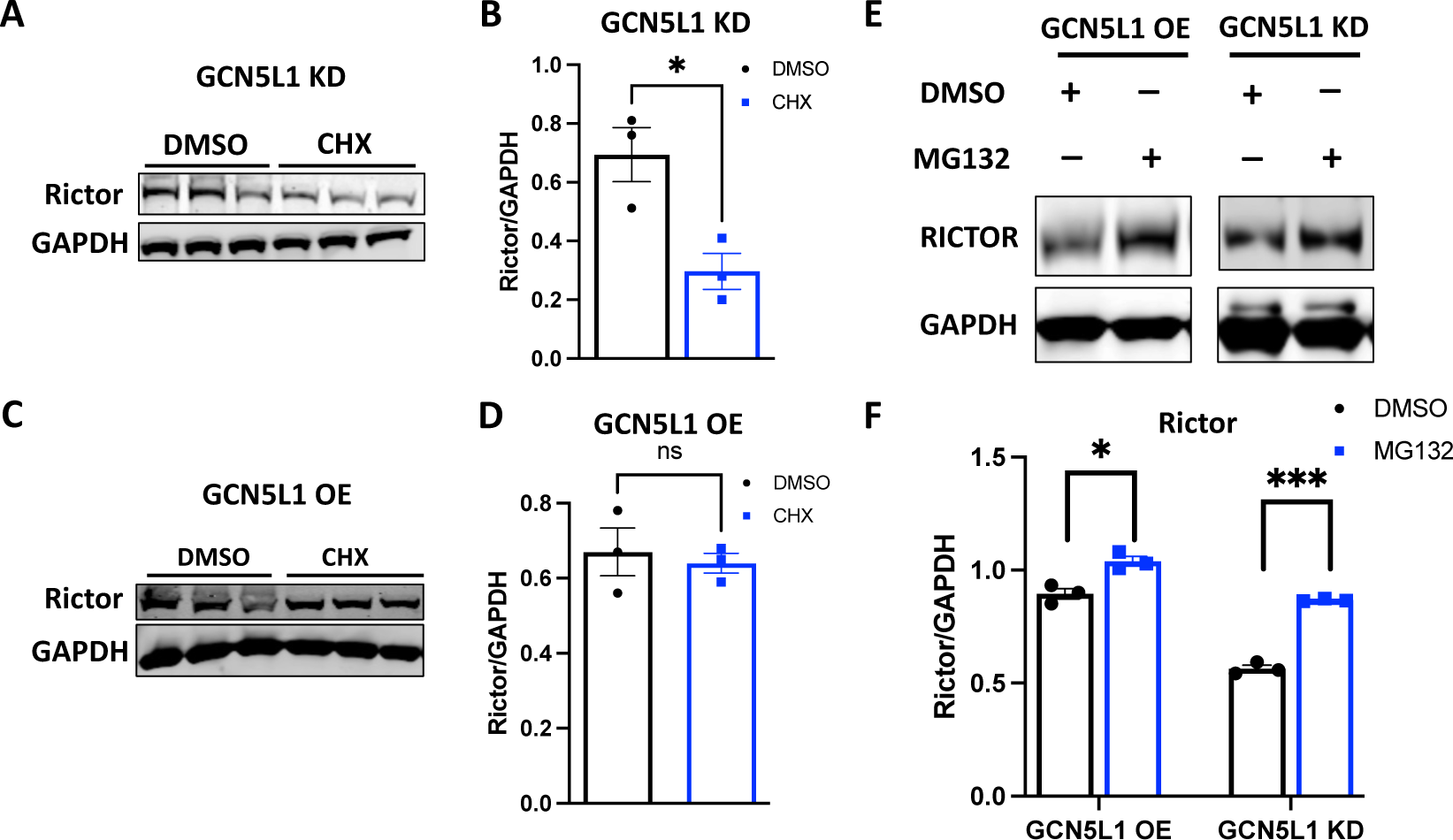
GCN5L1 regulates Rictor protein abundance in AC16 cells by decreasing proteasomal degradation. **A-B** Rictor protein levels of GCN5L1 KD AC16 cells after CHX treatment. **C-D** Rictor protein levels of GCN5L1 OE AC16 cells after CHX treatment. **E-F** Rictor protein levels in GCN5L1 KD and OE AC16 cells after MG132 treatment. Data were expressed as Mean±SEM, n=3, *p>0.05 vs DMSO ****p>0.001vs DMSO.

### Hypoxia-stimulated Rictor degradation is blocked by GCN5L1 acetyltransferase activity

Pathological circumstances alter protein levels without changing mRNA expression levels (Maejima, 2020). Our earlier experiments showed a significant decrease in Rictor protein abundance in AC16 cells when they underwent hypoxic stress, and we next attempted to understand the mechanisms underlying this change. Our previous experiment showed that GCN5L1 overexpression could inhibit Rictor protein degradation at the basal level by promoting Rictor acetylation. We therefore examined the role of GCN5L1 on hypoxia-induced Rictor degradation, and analyzed Rictor protein levels in GCN5L1 KD and OE AC16 cells after hypoxic injury. We witnessed a rapid degradation of Rictor protein in GCN5L1 KD cells, while GCN5L1 OE inhibited Rictor degradation (Fig. 6 A-B, E-F). As hypoxia creates nutrient-deprived conditions, we hypothesized that hypoxia should decrease Rictor acetylation levels in AC16 cells, and therefore quantified Ac-Rictor protein levels under normoxia and hypoxia conditions. We observed a significant decrease in Ac-Rictor levels when control or GCN5L1 KD AC16 cells underwent hypoxia stress (Fig. 6 C-D). Conversely, we found no change in Ac-Rictor levels in GCN5L1 OE AC16 cells (Fig 6 G-H). Based on these findings, we conclude that hypoxia results in decreased Rictor acetylation which promotes it proteasomal degradation, which can be blocked by increased GCN5L1 acetyltransferase activity.

**Figure 6.**
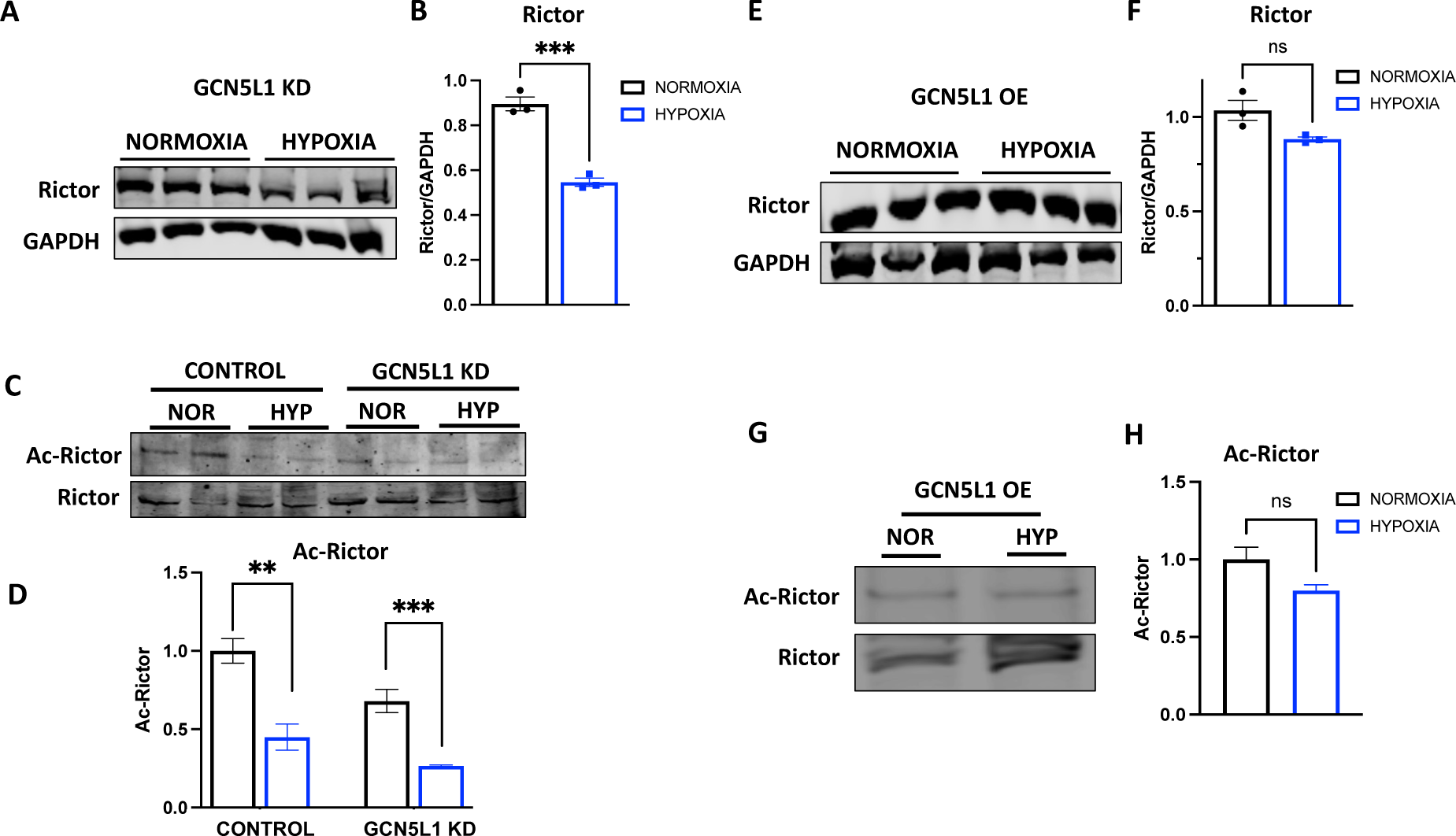
GCN5L1 activity reduces Rictor degradation after hypoxic injury in AC16 cells. **A-B** Rictor protein levels in GCN5L1 KD AC16 cells after hypoxia stress. **C-D** Ac-Rictor protein levels in control and GCN5L1 KD AC16 cells after hypoxia stress. **E-F** Rictor protein levels in GCN5L1 OE AC16 cells after hypoxia stress. **G-H** Ac-Rictor protein levels in GCN5L1 OE AC16 cells after hypoxia stress. Data were expressed as Mean±SEM, n=3, **p>0.01 vs Normoxia treated AC16 cells, ***p>0.001 vs Normoxia treated AC16 cells.

### Knockdown of GCN5L1 in AC16 cells restricts cytoprotective Akt/mTORC2 signaling via Rictor downregulation

Acetylation of Rictor promotes cancer cell survival via the attenuation of apoptosis (Masui et al., 2015). Cytoprotective signaling is essential to for the survival of any cell under stress, and it is crucial for non-regenerative tissues like cardiomyocytes, neurons, and skeletal muscle cells (He et al., 2022, Masmoudi-Kouki et al., 2020). Pathological conditions (e.g. hypoxia, injury, nutrient starvation, etc.) impair cytoprotective signaling of any cell/organ (Manning et al., 2019b). Rictor is a crucial component of mTORC2-mediated cell survival signaling, and our studies above demonstrate that Rictor protein levels decrease during hypoxia, which is further exacerbated by loss of GCN5L1 (Fig. 1 and Fig. 6). We therefore studied the Akt/mTORC2 cytoprotective signaling pathway in control and GCN5L1 KD AC16 cells, by measuring Rictor, Akt, p-Akt (Ser473), mTOR, and p-mTOR (Ser2448) protein levels. Activation of cytoprotective signaling is typically shown by increased levels of p-Akt (Ser473) and p-mTOR (Ser2448) (Manning et al., 2019b, Chadha and Meador-Woodruff, 2020). We observed a significant decrease in Rictor protein levels in control AC16 cells and GCN5L1 KD AC16 cells during hypoxic stress, with a greater decrease in Rictor protein evident in GCN5L1 KD AC16 cells when compared to control AC16 cells (Fig. 7 A-B). Cytoprotective p-Akt (Ser473) levels were significantly increased in control AC16 cells during hypoxic injury, and there were no significant changes in mTOR or p-mTOR levels following hypoxia (Fig. 7 A-B). In contrast, GCN5L1 KD cells showed a much lower increase in p-Akt (Ser473) levels, and displayed a decrease in mTOR and p-mTOR (Ser2448) levels in hypoxia (Fig 7 A-B). Combined, these data suggest that the magnitude of Rictor protein loss during ischemic injury determines the extent of cytoprotective signaling through the Akt/mTORC2 pathway, and that cytoprotective signaling is greatly attenuated in cells lacking GCN5L1.

**Figure 7.**
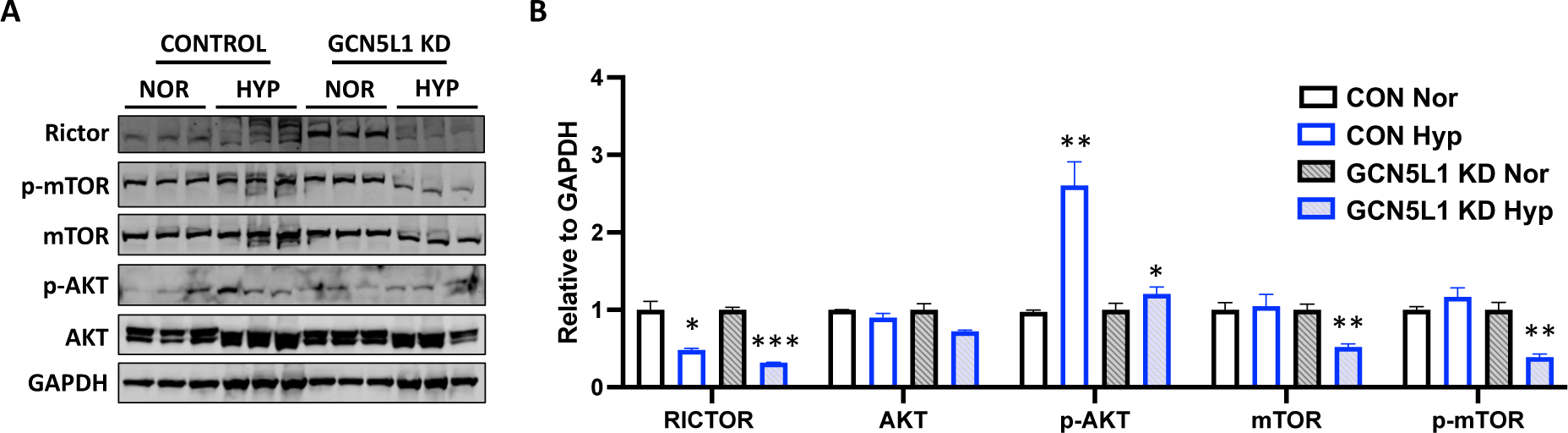
Cytoprotective Akt/mTORC2 signaling is attenuated in GCN5L1 knockdown cells. **A-B** mTORC2 mediated cytoprotective signaling proteins of control and GCN5L1 KD AC16 cells. Data were expressed as Mean±SEM, n=3, *p>0.01 vs Normoxia treated AC16 cells, **p>0.01 vs Normoxia treated AC16 cells, ****p>0.0005 vs Normoxia treated AC16 cells.

## Discussion

Ischemic heart disease is one of the major causes of death worldwide (Khan et al., 2020). A myriad of pathological mechanisms (e.g. Inflammation, apoptosis, fibrosis, etc.) are responsible for ischemic disease severity and mortality (Severino et al., 2020). Augmenting cytoprotective signaling, either through the prevention of cardiomyocyte apoptosis, or via cardiac regeneration, can be a potential therapeutic target to treat ischemic heart disease (Schirone et al., 2022). Rictor (RAPTOR Independent Companion of MTOR Complex 2) is crucial for mTORC2-mediated cytoprotective signaling (Zhao et al., 2014). The current study demonstrates that GCN5L1 regulates Rictor protein levels through modulation of Rictor post-translational modifications, and identified its role in cytoprotective signaling in AC16 cells during hypoxia stress. We initially measured the Rictor protein levels in nonfailing heart (NFH) and ischemic heart disease (IHD) patient cardiac tissue samples, and found that Rictor protein levels were significantly lowered in IHD cardiac tissue samples compared to NFH samples (Fig. 1). In both control AC16 cells subject to hypoxia, and GCN5L1 KD cells at baseline, we consistently found that Rictor protein levels were decreased in the absence of changes in Rictor mRNA expression. The abundance of a given protein can be regulated at two levels post-transcription, and involves either changes in mRNA translational activity/protein synthesis, or post-translational changes in protein degradation pathways (e.g.., lysosomal protein degradation, proteasomal degradation pathway) (Park et al., 2020, Ma et al., 2011, Li et al., 2022). We inhibited both pathways in control AC16 cells with CHX (mRNA translation inhibitor) and MG132 (proteasomal inhibitor), and measured Rictor protein levels. We observed that both pathways influence Rictor protein levels in AC16 cells, but inhibition of the proteasomal protein degradation pathway by MG132 predominantly leads to accumulation of Rictor protein levels. We therefore conclude that turnover of Rictor in response to cellular stress if the main mechanism controlling its abundance in cardiac cells.

While GCN5L1 is best known for its mitochondrial acetyltransferase activity (Scott et al., 2018, Zhang et al., 2023a), recently several of its non-mitochondrial targets have also been identified (e.g. Rictor and α-tubulin) (Wu et al., 2018, Manning et al., 2019a). We quantified Ac-Rictor levels in control and GCN5L1 KD AC16 cells, and found significantly lower Ac-Rictor levels in GCN5L1 KD cells (Fig 2 F-G). From this, we hypothesized that the GCN5L1 prevents Rictor degradation by promoting its acetylation. To further confirm the role of GCN5L1 on Rictor protein stability, we examined Rictor protein levels in GCN5L1 OE AC16 cells, and found that increased Rictor protein levels in GCN5L1 OE cells corresponded with elevated Ac-Rictor levels (Fig. 3). We therefore conclude that the abundance of acetyl modifications of Rictor are a major contributor to its stability in both normal and stress conditions.

Mechanistically, we propose that acetylation of Rictor by GCN5L1 protects the mature protein from proteasomal degradation. Loss of GCN5L1 expression in ischemic injury leads to a decrease in Rictor acetylation, which promotes its degradation. Several previous studies have shown that acetylation of lysine residues antagonizes ubiquitin-dependent degradation pathways, by blocking access of ubiquitin ligases to lysine residues. This prevents the addition of ubiquitin chains required for the initiation of proteasomal degradation (Li et al., 2012, Gronroos et al., 2002). As such, we hypothesize that GCN5L1-mediated acetylation of Rictor is important to maintain cytoprotective signaling through the Akt/mTORC2 signaling pathway, by maintaining Rictor protein abundance. Future work will determine whether supporting this pathway, and preventing Rictor deacetylation and degradation in ischemia, is a new potential therapeutic approach.

## Conclusion

The current research study is for the first time, reporting that ischemic stress decreases Rictor protein levels in the absence of changes to Rictor mRNA expression. GCN5L1 prevents Rictor degradation, promotes Rictor stability during hypoxic injury, and maintains cytoprotective signaling during hypoxia stress in AC16 cells.

### Limitations and Future Directions

We did not identify the acetylation sites of the Rictor protein that are modified by GCN5L1. In our future studies, we will identify the target residues on Rictor protein of GCN5L1, and examine whether modulation of these sites through acetyl-mimetic modifications is cytoprotective. As protein deacetylation makes cells susceptible to hypoxic stress, future studies can also examine whether protein deacetylase (HDAC) inhibitors are protective.

## Supplemental Figure

**Figure S1.**
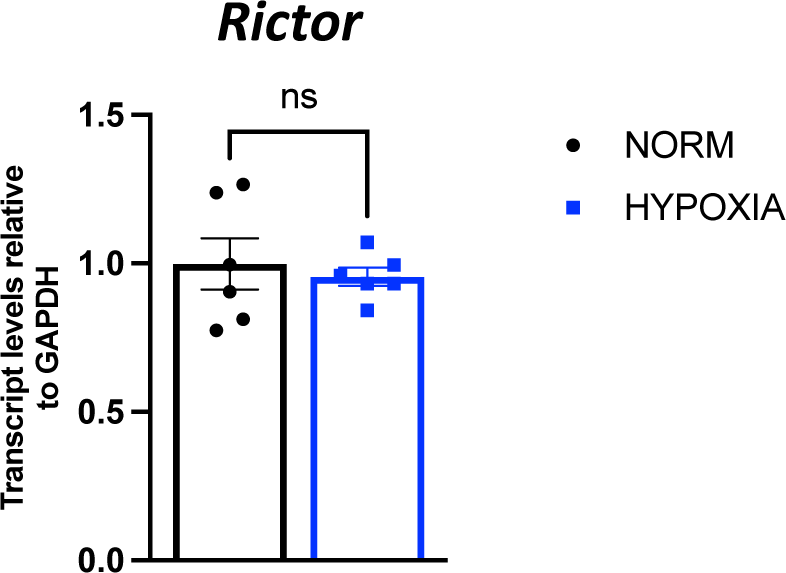
*In vitro* hypoxia does not affect Rictor mRNA expression in control AC16 cells. **A** Rictor mRNA expression levels in control AC16 cells after hypoxia stress. Data were expressed as Mean±SEM, n=5.

